# TreeArchTraits: an R package to analyse the architectural traits of trees using TLS data

**DOI:** 10.1101/2023.09.29.560266

**Authors:** Nicola Puletti, Simone Innocenti, Matteo Guasti, Cesar Alvites

## Abstract

1

The architecture of trees is significantly influenced by their interactions, directly affecting their functioning and structural development. Different tree architectural indicators (TAT) have evolved, including these interactions’ measurements.
Recent advances in Terrestrial Laser Scanning data collection make measuring the three-dimensional characteristics of trees more more efficient.
The R package `TreeArchTraits` facilitates the processing of three-dimensional tree characteristics obtained through Terrestrial Laser Scanning, enabling the computation of various indices.
A set of trees belonging to different forest conditions was used to demonstrate the `TreeArchTraits` potential for characterizing tree architectures.

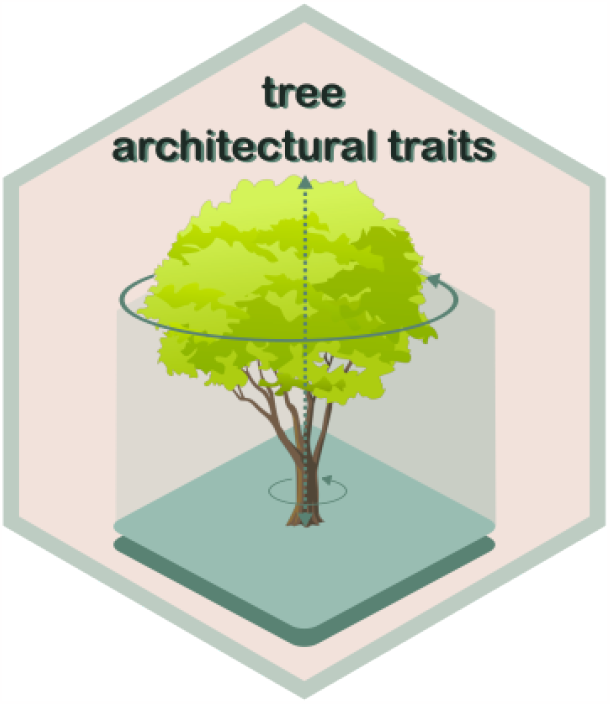

## 2 Background

The morphological architecture of trees is strongly affected by both their functional traits (physiological and/or phenological) and forest management practices. Therefore, analyzing how trees develop can enhance our understanding of the forest structure (Tomlinson, P. B. 1983).

Some authors (Dorji et al. 2021) have recently pointed out the importance of studying tree structure and form for diverse research fields, such as phylogeny and taxonomy, ecosystem modelling, tree physiology, and crucial for remote sensing of canopy landscapes, tree wind damage studies, carbon stock calculation for climate change mitigation schemes, as well as metabolic scaling theory (Malhi et al. 2018). Thus, knowledge of the individual architecture of trees is important but is limited today due to measurement complexity.

Compared to traditional methods, Terrestrial Laser Scanning (TLS) provides massive amounts of point cloud data to quantify three-dimensional attributes of whole forests and single trees.

In the package `TreeArchTraits` we collected a set of first architectural traits of trees that can be derived using TLS.

## 3 Package overview

`TreeArchTraits` has been developed as an R (R Core Team 2023) package. Currently, the most up-to-date version of the package can be downloaded free of charge from the GitLab development repository of the project (https://gitlab.com/Puletti/tree_arch_traits).

The functions available in the `TreeArchTraits` package are:

- `TraditionalTreeParam;sss`
- `StemParam;`
- `CrownParam;`
- `BoxDim;`
- `BoxDim_adj`.

An additional function to import *las* or *laz* files as a `data.table (las2datatable)` is also available. This function automatically converts the .las/.laz files into data.table, which is used for computing the five `TreeArchTraits` functions.

### 3.1 DBH and Total Tree Height estimation

The `TraditionalTreeParam` estimates the diameter-at-breast height and total Tree height for segmented tree point clouds (i.e. isolated and cleaned). The only argument required is this single tree object, imported as a `data.table` that can be derived using the `las2datatable` function.

### 3.2 Stem and crown parameters calculation

The StemParam function estimates a set of architectural traits from a TLS point cloud of a stem, returning a list of four elements: (1) the whole stem `data.frame;` (2) the DBH-centroid coordinates; (3) the crown-base-centroid `data.frame;` (2) the DBH-centroid coordinates; (3) the crown-base-centroid `data.frame`. The last list-element reports (4) the Stem Volume and some stem traits namely ‘Lean’, ‘Sweep’, and ‘Uprighgtness’, as from previous studies (Juchheim et al. 2017).

The function `CrownParam` estimates a set of crown architectural traits from a TLS point cloud of a crown (leaf-off conditions). The functions returns a list of 2 elements: (1) the whole crown `data.frame` and (2) the surface area (`CSA`, in *m*^2^), the crown volume (`Cvol`, in *m*^3^), and an estimates of branch volume (`Bvol`, in *m*^3^) as a first list of crown attributes. As for `StemParam`, the crown volume is computed using a convex hull approach in slices.

### 3.3 Box dimension

Box-dimension parameter (*Db*) is an integrative measure of tree architecture firstly suggested by Mandelbrot (Mandelbrot 1982). Considering the TLS technology potential for a 3D detailed analysis, *Db* was recently re-introduced by some authors (Seidel et al. 2019; Heidenreich and Seidel 2022; Neudam, Annighöfer, and Seidel 2022). *Db* is a measure sensitive to both outer shape and internal structure of a tree. It is also scale independent and hence useful when trees of different sizes must be compared.

*Db* is determined by evaluating how many virtual boxes one needs to enclose all points of the 3D tree and how the number of boxes changes with the ratio of the box size to the original box size. *Db* is then the slope of the fitted straight line through the scatterplot of the logarithm of the number of boxes needed over the inverse of the logarithm of the used box-resolution. For more details on *Db* calculations, please refer to specific experiments.

In this package, `BoxDims` is a first version function for the estimates the *Db*. The arguments are:

- a `data.table (pc_tree)` obtained by the `las2datatable` function, and
- the number of points within a voxel to define it as “vegetation” (`pvt`).

The output of this function is a `list` containing the estimated box-dimension as a measure of overall tree complexity, the box-dimension_intercept as a measure of tree size, and the whole `data.frame` as last element.

`BoxDim_adj` is new version of this function, closely related to computations performed by (Seidel et al. 2019). The main difference with `BoxDim` is that the baricenter of the tree-point-cloud and the baricenter of the first main biggest voxel were coincident. The input is only the `data.table (pc_tree)` representing the tree point-cloud data, while the output is a `data.frame` with: (i) the number of voxel containing the whole tree and (ii) the corresponding edge length. The output of this function can be directly used to perform *Db* computation and graphics (see Figure 1).

**Figure 1.**
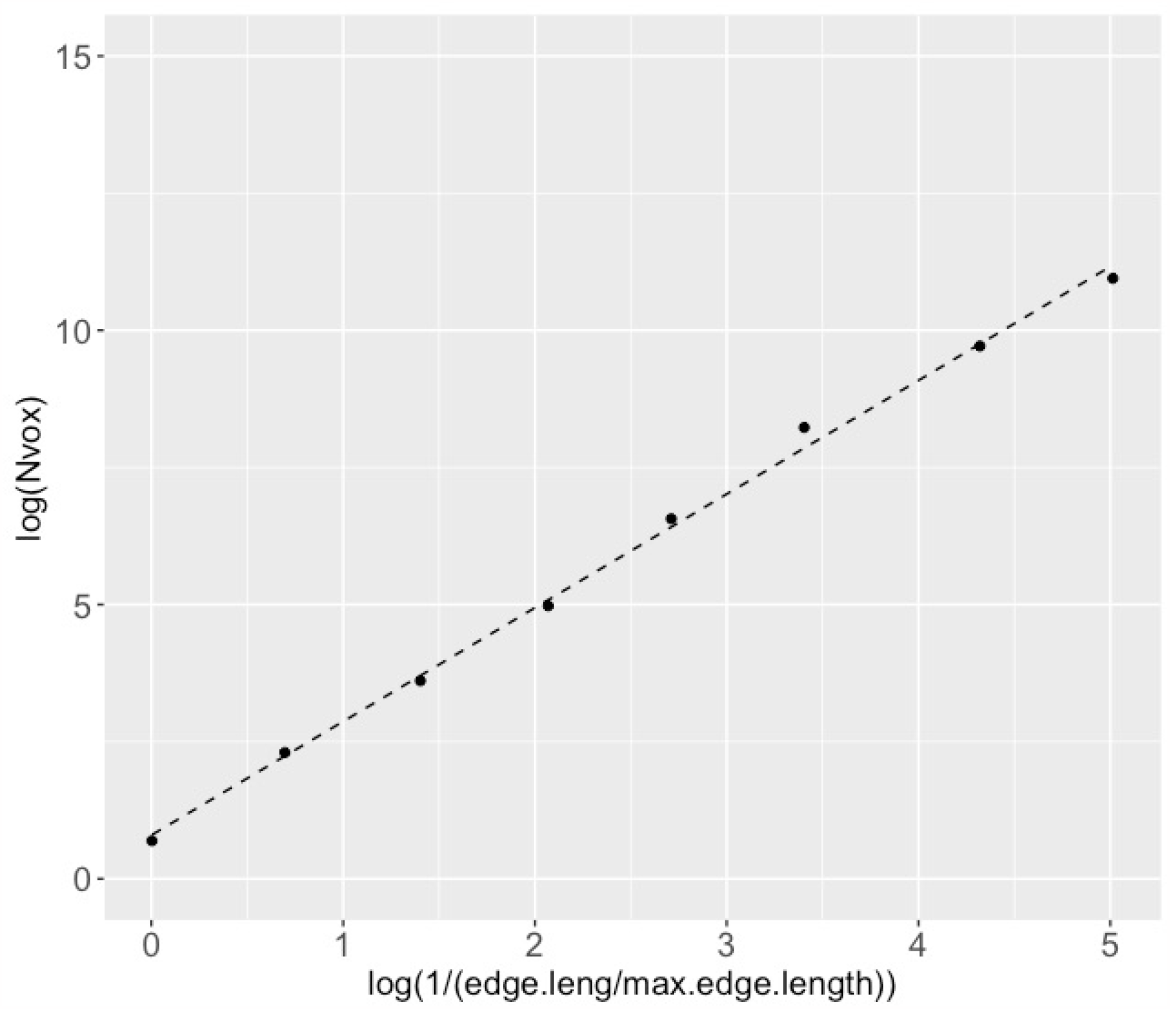
Exemplary log–log plot of the number of boxes [Nvox] over the inverse of the box size for the point cloud of a Chestnut tree (Castanea Sativa M.). For this tree Db is equal to 2.07, while the “tree size index” (i.e. the intercept of the liner regression line) is equal to 0.802.

## 4 Example application

### 4.1 Data collection and point cloud pre-processing

The examples presented in the following paragraphs use TLS data collected in field surveys from different projects in Italy. The entire dataset is stored in Zenodo https://zenodo.org/record/7285178#.ZDpN4ezP08E. All the trees were manually segmented by an expert operator using Trimble Realworks®software (Figure 2). These point-clouds can be used for both `TraditionalTreeParam, BoxDim`, and `BoxDim_adj` functions.

**Figure 2.**
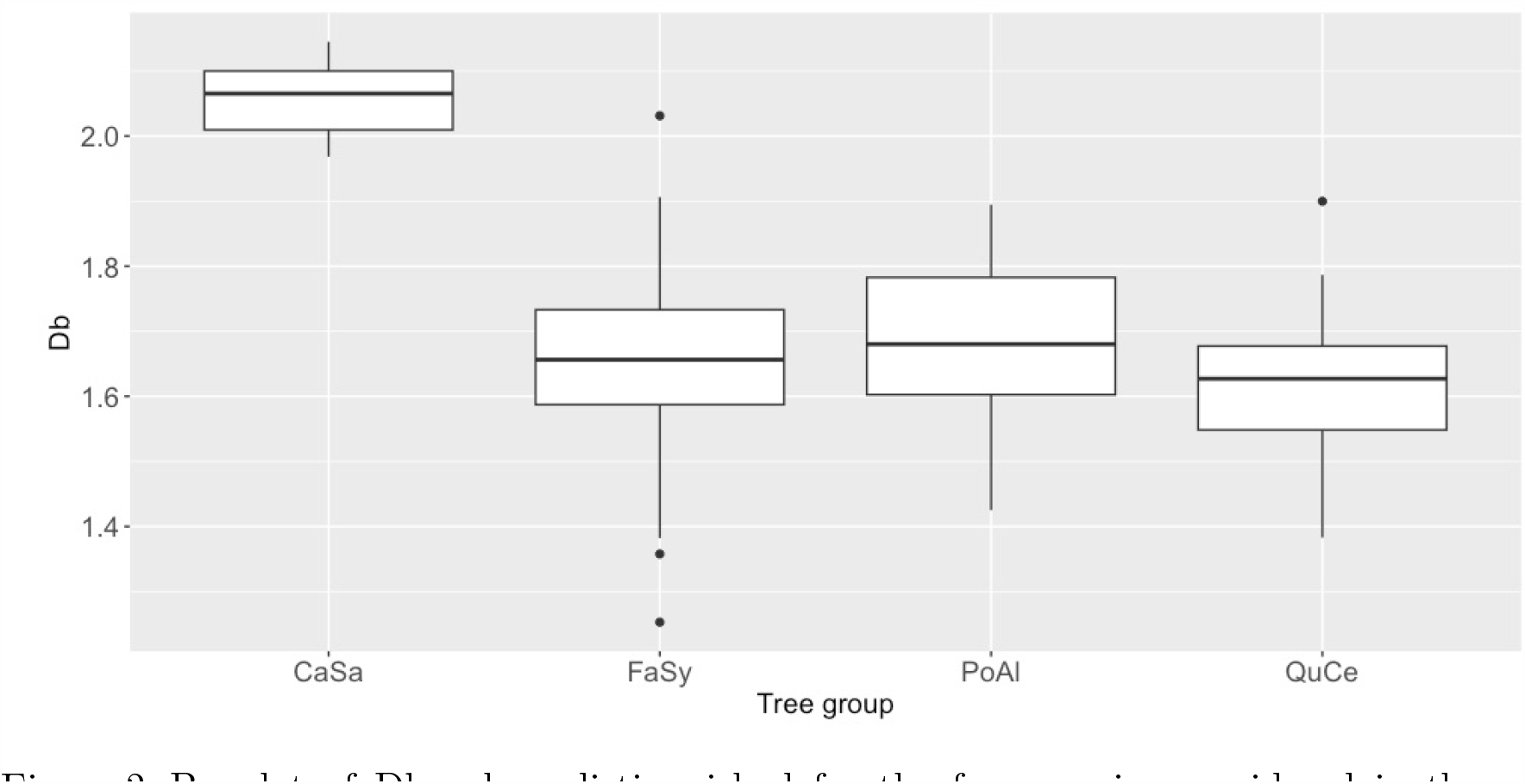
Boxplot of Db values distinguished for the four species considered in the case studies presented in this paper. CaSa (*Castanea sativa*), FaSy (*Fagus sylvatica*), PoAl (*Populus alba*) and QuCe (*Quercus cerris*).

**Figure 3.**
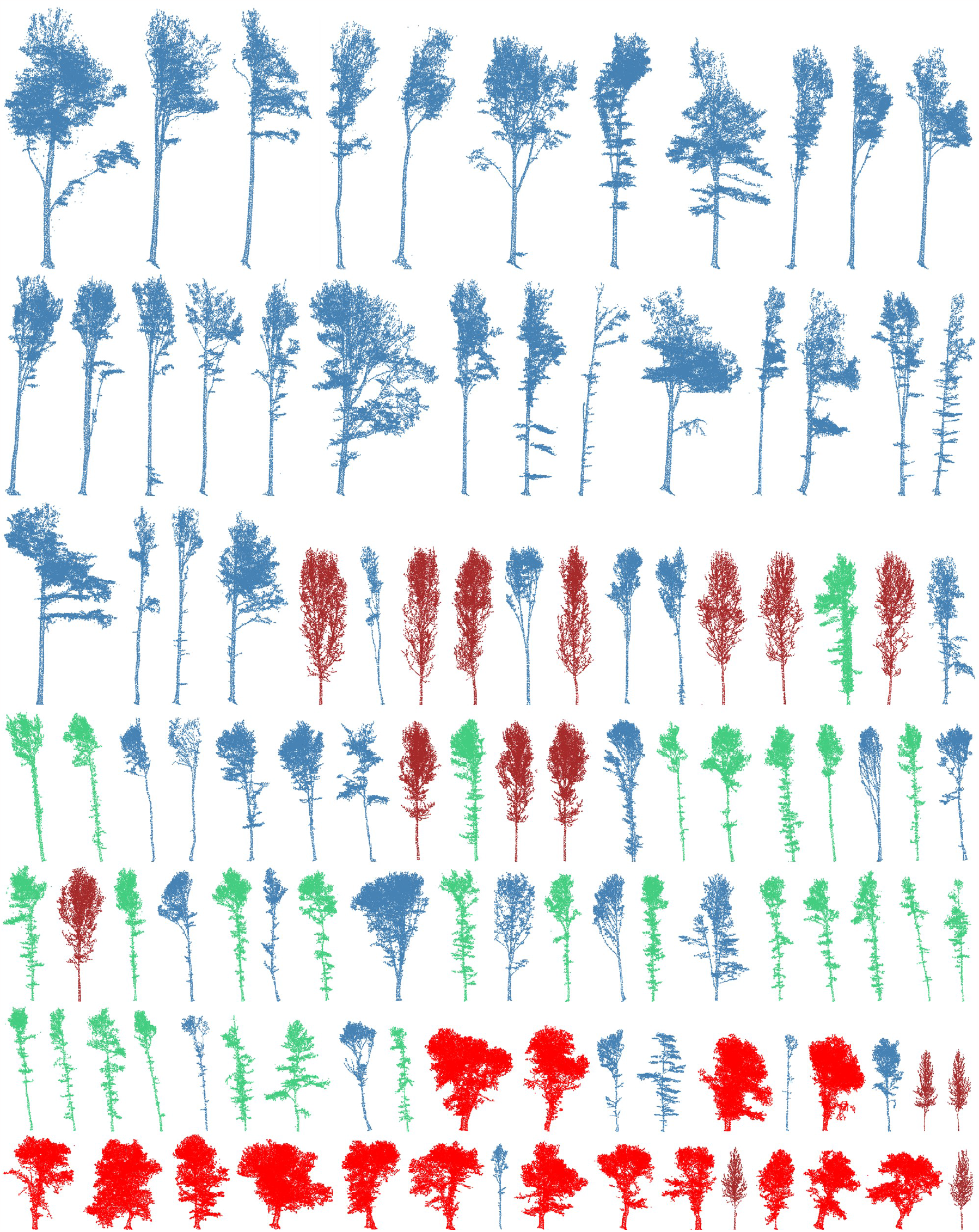
The total list of trees arranged from the tallest to the lowest. Steelblue colour is referred to Beech trees, the Oak trees are green, the Chestnut trees are red, and the Poplar trees are brown.

To analyse stem and crown architectural traits, using respectively `StemParam` and `CrownParam` functions, each tree was manually sub-segmented into two files, one for crown and one for stem.

### 4.2 Architectural traits: case studies

In the case studies presented in this paper, we want to focus particularly on box-dimension parameter (*Db*). We compare different forest conditions listed here below.

1. <u>Broadleaved tree species.</u> It is a dataset of 48 trees: 12 Chestnuts (*Castanea sativa, Casa*), 12 Beeches (*Fagus sylvatica, FaSy*), 12 Poplars (*Populus alba, PoAl*), and 12 Oaks (*Quercus cerris, QuCe*) (Figure 2). With this subset we want to highlights Db potential in characterizing tree-traits of quite similar species;
2. <u>Forest management (same species).</u> 2a) Beech (*Fagus sylvatica*): 25 trees from Sasso Fratino, and 25 trees from traditional forest management (Figure 4 on the left); 2b) Turkey oak (*Quercus cerris*): 24 trees from three different forest management (Figure 4 on the right);
3. <u>Age classes (same species)</u> Here we consider the same species (Poplar plantation, *Populus alba*) to analyse changes in time (*Populus alba, PoAl*) (Figure 5).

**Figure 4.**
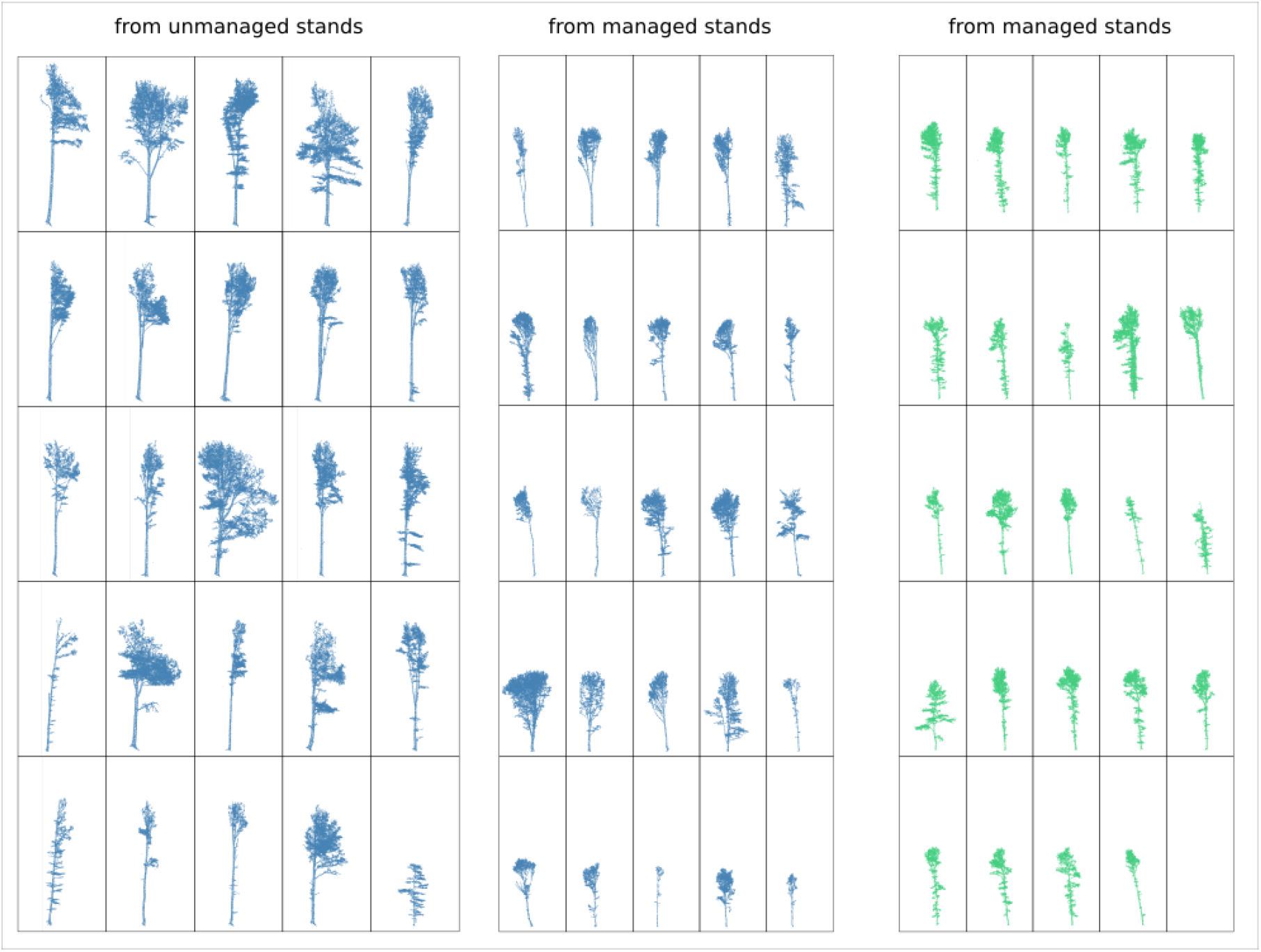
List of Beech trees (blue color) from unmanaged stands (Foreste Casentinesi National Park) and from managed stands used in this case study. In green the list of Oak trees from managed stands.

**Figure 5.**
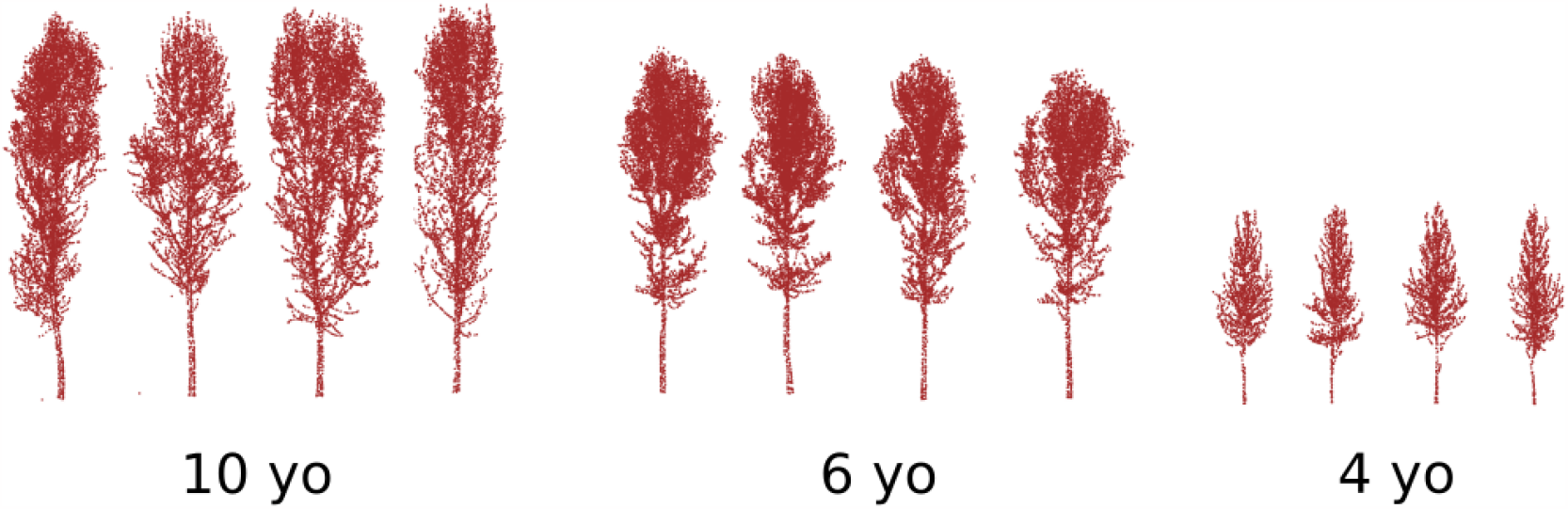
Point clouds for poplar trees collected in three different age classes: 4, 6, and 10 years old.

#### 4.2.1 Differences in broadleaved tree species

The question here is if *Db* is able to distinguish architecture of similar species. We used data on broadleaves forests, collected in four sites located in Northern Italy. On average, the fractal dimension of trees, which can be estimated by the *Db*, is smaller than two for trees growing from mid to high competition (like beeches, oaks, and poplars). Instead, *Db* is higher than 2 for chestnuts, because they are trees grew in the open.

| Species group | N | median of $Db$ | sd of $Db$ |
| --- | --- | --- | --- |
| FaSy_F | 25 | 1.69 | 0.129 |
| FaSy_T | 25 | 1.62 | 0.163 |
| QuCe_Bb | 8 | 1.65 | 0.104 |
| QuCe_Bs | 8 | 1.66 | 0.162 |
| QuCe_Co | 8 | 1.64 | 0.069 |

### 4.2.2 Differences in forest management

Forest management is a key driver for tree architecture. Data of unmaneged Beech trees were collected within the protected forest area of Sasso Fratino, Foreste Casentinesi National Park, northern Apennines, Italy. Data of managed Beech trees were collected in the ‘Alpe di Catenaia’ mountain forest, Tuscany. The climate at both the study sites is temperate, with warm, dry summers and cold, rainy winters.

| Species group | N | median of $Db$ | sd of $Db$ |
| --- | --- | --- | --- |
| CaSa | 12 | 2.05 | 0.056 |
| FaSy | 12 | 1.61 | 0.095 |
| PoAl | 12 | 1.67 | 0.149 |
| QuCe | 12 | 1.63 | 0.091 |

#### 4.2.3 Differences in age classes

Competition between trees increases with age, particularly from the moment the crown of a tree touches the crowns of their surrounding trees. To evaluate the potential of the package in analysing architectural traits of trees that compete in time, we used multitemporal-data of poplar plantation. Such trees were planted using 6 m by 7 m arrangement.

| Age | N | median $Db$ | sd $Db$ |
| --- | --- | --- | --- |
| 4 years | 4 | 1.66 | 0.080 |
| 6 years | 4 | 1.73 | 0.128 |
| 8 years | 3 | 1.77 | 0.207 |
| 10 years | 4 | 1.64 | 0.178 |

## Acknowledgments

This research was funded by the Italian Ministry of Agriculture, Food, and Forestry Policies (MiPAAF), subproject “Precision Forestry” of the project AgriDigit program-DM, funding number 36503.7305.2018 of 2 December 2018).

